# Atomistic Simulations and Deep Mutational Scanning of Protein Stability and Binding Interactions in the SARS-CoV-2 Spike Protein Complexes with Nanobodies: Molecular Determinants of Mutational Escape Mechanisms

**DOI:** 10.1101/2021.07.07.451538

**Authors:** Gennady M. Verkhivker, Steve Agajanian, Deniz Yasar Oztas, Grace Gupta

## Abstract

Structural and biochemical studies have recently revealed a range of rationally engineered nanobodies with efficient neutralizing capacity against SARS-CoV-2 virus and resilience against mutational escape. In this work, we combined atomistic simulations and conformational dynamics analysis with the ensemble-based mutational profiling of binding interactions for a diverse panel of SARS-CoV-2 spike complexes with nanobodies. Using this computational toolkit we identified dynamic signatures and binding affinity fingerprints for the SARS-CoV-2 spike protein complexes with nanobodies Nb6 and Nb20, VHH E, a pair combination VHH E+U, a biparatopic nanobody VHH VE, and a combination of CC12.3 antibody and VHH V/W nanobodies. Through ensemble-based deep mutational profiling of stability and binding affinities, we identify critical hotspots and characterize molecular mechanisms of SARS-CoV-2 spike protein binding with single ultra-potent nanobodies, nanobody cocktails and biparatopic nanobodies. By quantifying dynamic and energetic determinants of the SARS-CoV-2 S binding with nanobodies, we also examine the effects of circulating variants and escaping mutations. We found that mutational escape mechanisms may be controlled through structurally and energetically adaptable binding hotspots located in the host receptor-accessible binding epitope that are dynamically coupled to the stability centers in the distant epitope targeted by VHH U/V/W nanobodies. The results of this study suggested a mechanism in which through cooperative dynamic changes, nanobody combinations and biparatopic nanobody can modulate the global protein response and induce the increased resilience to common escape mutants.

## Introduction

SARS-CoV-2 infection is initiated when the viral spike (S) glycoprotein penetrates to the host cell receptor ACE2 leading to the entry of S protein into host cells and membrane fusion.^1–3^ The SARS-CoV-2 S protein consists of two subunits S1 and S2, where the subunit S1 includes an N-terminal domain (NTD) and the receptor-binding domain (RBD).^4, 5^ The rapidly growing body of cryo-EM structures of the SARS-CoV-2 S proteins revealed distinct conformational arrangements of the S protein trimers in the prefusion form that display a dynamic equilibrium between the closed (“RBD-down”) and the receptor-accessible open (“RBD-up”) form required for the S protein fusion to the viral membrane.^6–18^ A series of conformational events associated with ACE2 binding include progression from a compact closed form of the SARS-CoV-2 S protein to the partially open states and ACE2-bound form priming the protein for fusion activation.^19^ Functional adaptability and conformational plasticity of SARS-CoV-2 S proteins allow for efficient modulation of selective phenotypic responses to binding of the host receptor ACE2 and the emerging spectrum of neutralizing antibodies.^20–28^ Some ultra-potent antibodies displayed high potency by binding to the NTD regions and allosterically interfering with the host receptor binding without directly interfering with ACE2 recognition.^28^ SARS-CoV-2 antibodies can be divided into several main classes, of which class I and class II antibodies represent the dominant group of structurally characterized monoclonal antibodies that target the receptor binding motif (RBM) region of the RBD overlap with the ACE2 epitope.^29–31^ Structural and biochemical studies characterized the binding epitopes and molecular mechanisms for a number of SARS-CoV-2 antibodies including REGN10933, B38, CB6, CC12.3 and BD-236 that compete for the RBD binding with the host receptor.^32–38^ Optimally designed antibody cocktails simultaneously targeting different binding epitopes on the SARS-CoV-2 RBD demonstrated excellent neutralization effects and improved resilience against mutational escape. Functional mapping of mutations in the SARS-CoV-2 S-RBD using deep mutational scanning showed that sites of antibody mutational escape are constrained by requirement of the protein stability and ACE2 binding, suggesting that escape-resistant antibody cocktails can limit the virus ability to acquire novel sites of immune escape in the RBD without compromising its binding to ACE2.^39–43^

Although functionally analogous to antibodies, nanobodies are much smaller and often more robust and thermostable. An ultra-potent synthetic nanobody Nb6 neutralizes SARS-CoV-2 by stabilizing the fully inactive down S conformation preventing binding with ACE2 receptor.^44^ Structure-guided design produced a high-affinity trivalent nanobody mNb6-tri that simultaneously binds to all three RBDs and inhibits the interactions with the host receptor by occupying the binding site and locking the S protein in the inactive, receptor-inaccessible state.^44^ Using a combination of size-exclusion chromatography, mass spectrometry, and structural modeling, a recent study revealed high-affinity nanobodies that target the RBD and efficiently neutralize SARS-CoV-2 by using several distinct and non-overlapping epitopes.^45^ This study also confirmed a dominant epitope targeted by Nb20 and Nb21 nanobodies that overlaps with the ACE2 binding site, showing that these nanobodies could competitively inhibit ACE2 binding and exploit structural mimicry w to facilitate conformational changes that prematurely convert spike into a post-fusion state suppressing viral fusion.^45^ Potent neutralizing nanobodies that resist circulating variants of SARS-CoV-2 by targeting novel epitopes were recently discovered.^46^ The reported cryo-EM structures for different classes of nanobodies suggested mechanisms of high-affinity and broadly neutralizing activity by exploiting epitopes that are shared with antibodies as well as novel epitopes that are unique to the nanobodies.^46^ Another diverse repertoire of high affinity nanobodies against SARS-CoV-2 S protein that are refractory to common escape mutants and showed synergistic neutralizing activity are characterized by proximal but non-overlapping epitopes suggesting that multimeric nanobody combinations can improve potency while minimizing susceptibility to escape mutations.^47^ These studies identified a group of common resistant mutations in the dynamic RBM region (F490S, E484K, Q493K, F490L, F486S, F486L, and Y508H) that evade many individual nanobodies.^47^ Structural versatility of nanobody combinations that can effectively insulate the S-RBD accessible regions suggested a mechanism of resistance to mutational escape in which combining two nanobodies can markedly reduce the number of allowed substitutions to confer resistance and thereby elevate the genetic barrier for escape.^47, 48^ Using human VH-phage library and protein engineering several unique VH binders were discovered that recognized two separate epitopes within the ACE2 binding interface with nanomolar affinity.^48^ Multivalent and bi-paratopic VH constructs showed markedly increased affinity and neutralization potency to the SARS-CoV-2 virus when compared to standalone VH domain.^48^ Using saturation mutagenesis of the RBD exposed residues combined with fluorescence activated cell sorting for mutant screening, escape mutants were identified for five nanobodies and were mostly mapped to periphery of the ACE2 binding site with K417, D420, Y421, F486, and Q493 emerging as notable hotspots.^49^

A wide range of rationally engineered nanobodies with efficient neutralizing capacity and resilience against mutational escape was recently unveiled that included the llama-derived nanobody VHH E bound to the ACE2-binding epitope and three alpaca-derived nanobodies VHHs U, V, and W that bind to a different cryptic RBD epitopee.^50–52^ Using X-ray crystallography and surface plasmon resonance-based binding competition this illuminating study showed that combinations of nanobodies targeting distinct epitopes can reduce the emergence of escape mutants that were resistant to individual nanobodies, while the biparatopic VHH EV and VE nanobodies with two antigen-binding sites were superior to pair combinations VHH E+U, E+V and E+W in preventing mutual escape.^50^ Using single-domain antibody library and PCR-based maturation, two closely related and highly potent nanobodies, H11-D4 and H11-H4 were reported that recognize the same epitope immediately adjacent to and partly overlapping with the ACE2 binding region.^53^ The crystal structures of these nanobodies bound to the S-RBD revealed binding to the same epitope, which partly overlaps with the ACE2 binding surface, explaining competitive inhibition of ACE2 interactions. These studies demonstrated that nanobodies may have potential clinical applications due to the increased neutralizing activity and robust protection against escape mutations of SARS CoV-2.

The global circulating mutants of the SARS-CoV-2 S protein discovered through epidemiological surveillance prompted a significant effort in the biomedical community. The B.1.1.7 variant of the SARS-CoV-2 featured N501Y mutation that can increase binding affinity with ACE2 while eliciting immune escape and reduced neutralization of RBD-targeting antibodies.^54–56^ A SARS-CoV-2 lineage (501Y.V2) is characterized by mutations K417N, E484K and N501Y that can also induce significant immune escape.^57–60^ The new lineage P.1 contains a constellation of lineage defining mutations, including E484K, K417T, and N501Y mutations.^61, 62^ Highly transmissible SARS-CoV-2 variants with the increased affinity for ACE2 were discovered in lineage B.1.617 that encodes for mutations L452R, T478K, E484Q, D614G and P681R as well as B.1.618 lineage with mutations Δ145-146, E484K and D614G.^63, 64^ The B.617.2 variant with L452R and T478K can induce a significant escape to neutralizing antibodies targeting the NTD and RBD, and polyclonal antibodies elicited by previous SARS-CoV-2 infection or vaccination. Nanobody and antibody mixtures targeting non-overlapping epitopes on the RBD have shown promise in preventing occurrence of resistance mutations. The reported high-affinity nanobodies and nanobody cocktails consisting of two noncompeting nanobodies potently neutralize both wild-type SARS-CoV-2 and the N501/ D614G variants.^65^ These studies suggested that nanobody mixtures and rationally engineered biparatopic nanobody constructs could offer a promising alternative to conventional monoclonal antibodies and may be advantageous for controlling a broad range of infectious variants while also suppressing the emergence of virus escape mutations.

Computer simulations and protein modeling played an important role in shaping up our understanding of the dynamics and function of SARS-CoV-2 glycoproteins.^66–76^ All-atom MD simulations of the full-length SARS-CoV-2 S glycoprotein embedded in the viral membrane, with a complete glycosylation profile were first reported by Amaro and colleagues, providing the unprecedented level of details and significant structural insights about functional S conformations.^69^ A “bottom-up” coarse-grained (CG) model of the SARS-CoV-2 virion integrated data from cryo-EM, x-ray crystallography, and computational predictions to build molecular models of structural SARS-CoV-2 proteins assemble a complete virion model.^70^ By providing valuable insights and establishing the blueprint for computational modeling, these studies paved the way for simulation-driven studies of SARS-CoV-2 spike proteins, also showing that conformational plasticity and the alterations of the SARS-CoV-2 spike glycosylation can synergistically modulate complex phenotypic responses to the host receptor and antibodies. Multi-microsecond MD simulations of a 4.1 million atom system containing a patch of viral membrane with four full-length, fully glycosylated and palmitoylated S proteins allowed for a complete mapping of generic antibody binding signatures and characterization of the antibody and vaccine epitopes.^71^ Our recent studies combined coarse-grained and atomistic MD simulations with coevolutionary analysis and network modeling to present evidence that the SARS-CoV-2 spike protein function as allosterically regulated machine that exploits plasticity of allosteric hotspots to fine-tune response to antibody binding.^77–82^ These studies showed that examining allosteric behavior of the SARS-CoV-2 pike proteins may be useful to uncover functional mechanisms and rationalize the growing body of diverse experimental data. A critical review of computational simulation studies of the SARS-CoV-2 S proteins highlighted the synergies between experiments and simulations, outlining directions for computational biology research in understanding mechanisms of COVID-19 protein targets.^83^

In this study, we combine atomistic simulations and conformational dynamics analysis with the ensemble-based mutational profiling of binding interactions for a diverse panel of SARS-CoV-2 S-RBD complexes with nanobodies. Using this computational toolkit we identified dynamic signatures and binding affinity fingerprints for the SARS-CoV-2 complexes with nanobodies Nb6 and Nb20, VHH E, a pair combination VHH E+U, a biparatopic nanobody VHH VE, and a combination of CC12.3 antibody and VHH V/W nanobodies. Through ensemble-based deep mutational profiling of stability and binding affinities, we identify critical hotspots and characterize molecular mechanisms of SARS-CoV-2 S binding with single ultra-potent nanobodies, nanobody cocktails and biparatopic nanobodies. By quantifying dynamic and energetic determinants of the SARS-CoV-2 S binding with nanobodies, we also examine the effects of circulating variants and escaping mutations. This study reveals that structural stability centers in the SARS-CoV-2 S-RBD are dynamically coupled to conformationally adaptable hotspots that control mutational escape. We show that through cooperative dynamic and energetic changes, nanobody combinations and biparatopic nanobody can modulate the global protein response and induce the increased resilience to common escape mutants.

## Results and Discussion

### Conformational Dynamics of the SARS-CoV-2 S RBD Complexes Reveals Nanobody-Induced Modulation of Spike Plasticity

We performed all-atom MD simulations of the SARS-CoV-2 S-RBD protein complexes with a panel of nanobodies (Figure 1) to examine how structural plasticity of the RBD regions can be modulated by binding and determine specific dynamic signatures induced by different classes of nanobodies targeting distinct binding epitopes. Overall, the results showed that nanobodies can induce a significant stabilization of the S-RBD in the protein core and RBM residues (Figure 2). A comparative analysis of the conformational flexibility profiles for complexes with Nb20, VHH E, and VHH VE revealed stabilization of the interacting regions that was especially pronounced in the complex with the biparatopic nanobody VHH VE (Figure 2A). The RBD core regions including α-helical segments of the RBD (residues 349-353, 405-410, and 416-423) showed only very small thermal fluctuations and remained stable. Furthermore, the conformational dynamics profiles for the nanobody complexes demonstrated stability of the central β strands (residues 354-363, 389-405, and 423-436) (Figure 2A). Of functional importance are anti-parallel β-sheets (β5 and β6) (residues 451-454 and 491-495) that connect the RBM region (residues 438-508) to the RBD core. These stable segments are involved in the binding interactions and are also proximal to sites of common mutational escape shared by many nanobodies such as L452, F486, and F490. At the same time, an appreciable level of flexibility was seen for the S-RBD complexes with Nb20 and VHH E in regions 359-372 and 380-390 that are distant from the RBM binding epitope (Figure 2A). Structural maps of the conformational dynamics profiles for the S-RBD complexes with Nb20 (Figure 2C) and VHH E (Figure 2D) illustrated the greater mobility of the RBM residues and plasticity of the binding epitope.

**Figure 1.**
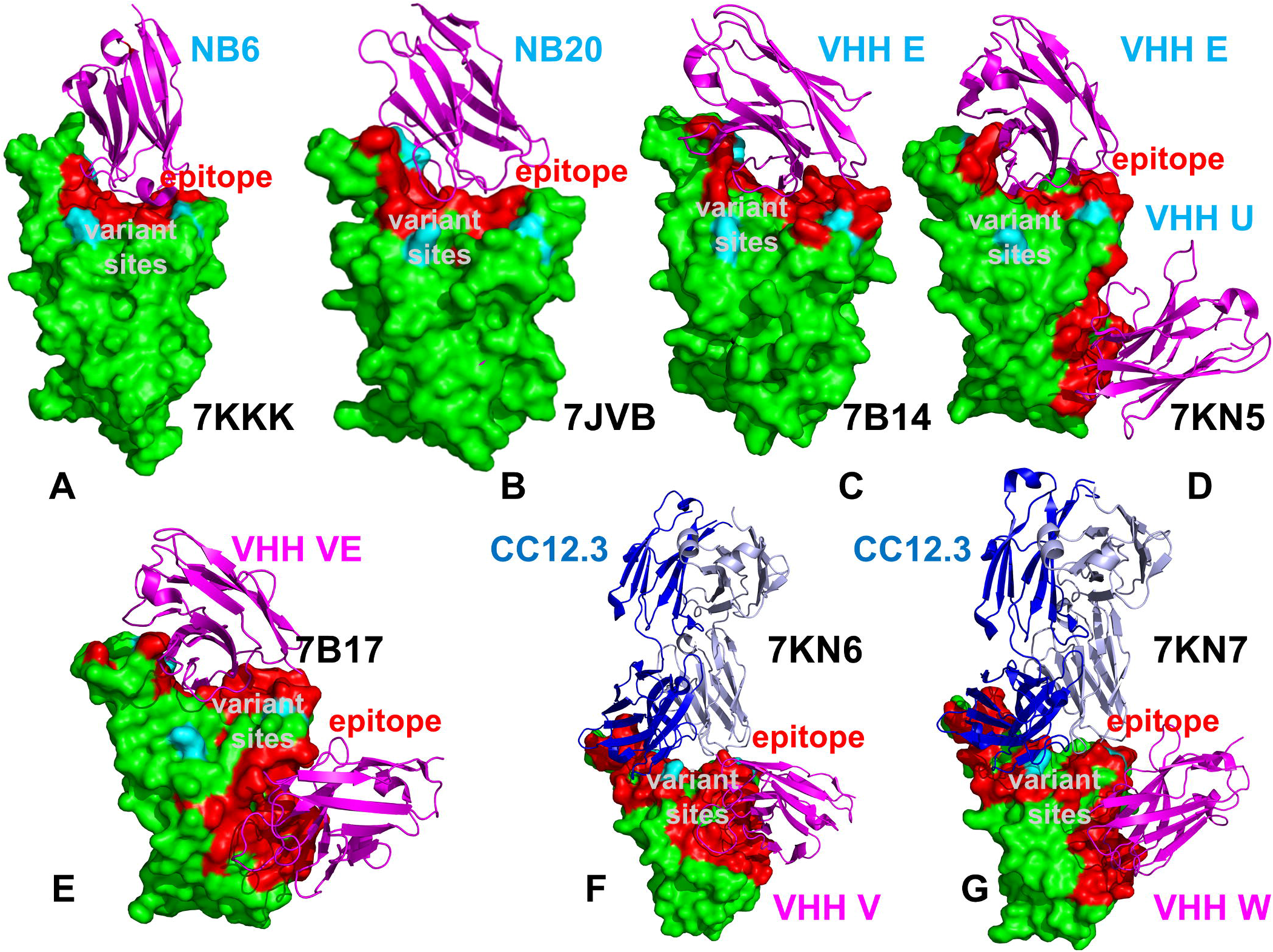
Crystal structures of the SARS-CoV-2 S trimer structures used in this study. The structures of SARS-CoV-2 S-RBD in the complex with Nb6 nanobody, pdb id 7KKK (A), complex with Nb20 nanobody, pdb id 7JVB (B), complex with VHH E nanobody, pdb id 7KN5 (C), a nanobody combination pair VHH E/ VHH U, pdb id 7KN5 (D), biparatopic nanobody VHH VE, pdb id 7B17 (E), a combination of CC12.3 antibody and VHH V nanobody, pdb id 7KN6 (F), and a combination of CC12.3 antibody and VHH W nanobody, pdb id 7KN7 (G). The S-RBD structures are shown in green surfaces. The binding epitope residues of the S-RBD bound structures are shown in red. The bound nanobody structures are shown in magenta ribbons and annotated. CC12.3 antibody on panels F and G is shown in ribbons with the heavy chain in blue and light chain in cyan colors. The sites K417, L452, E484 and N501 subjected to circulating mutational variants are shown in cyan surface and annotated.

**Figure 2.**
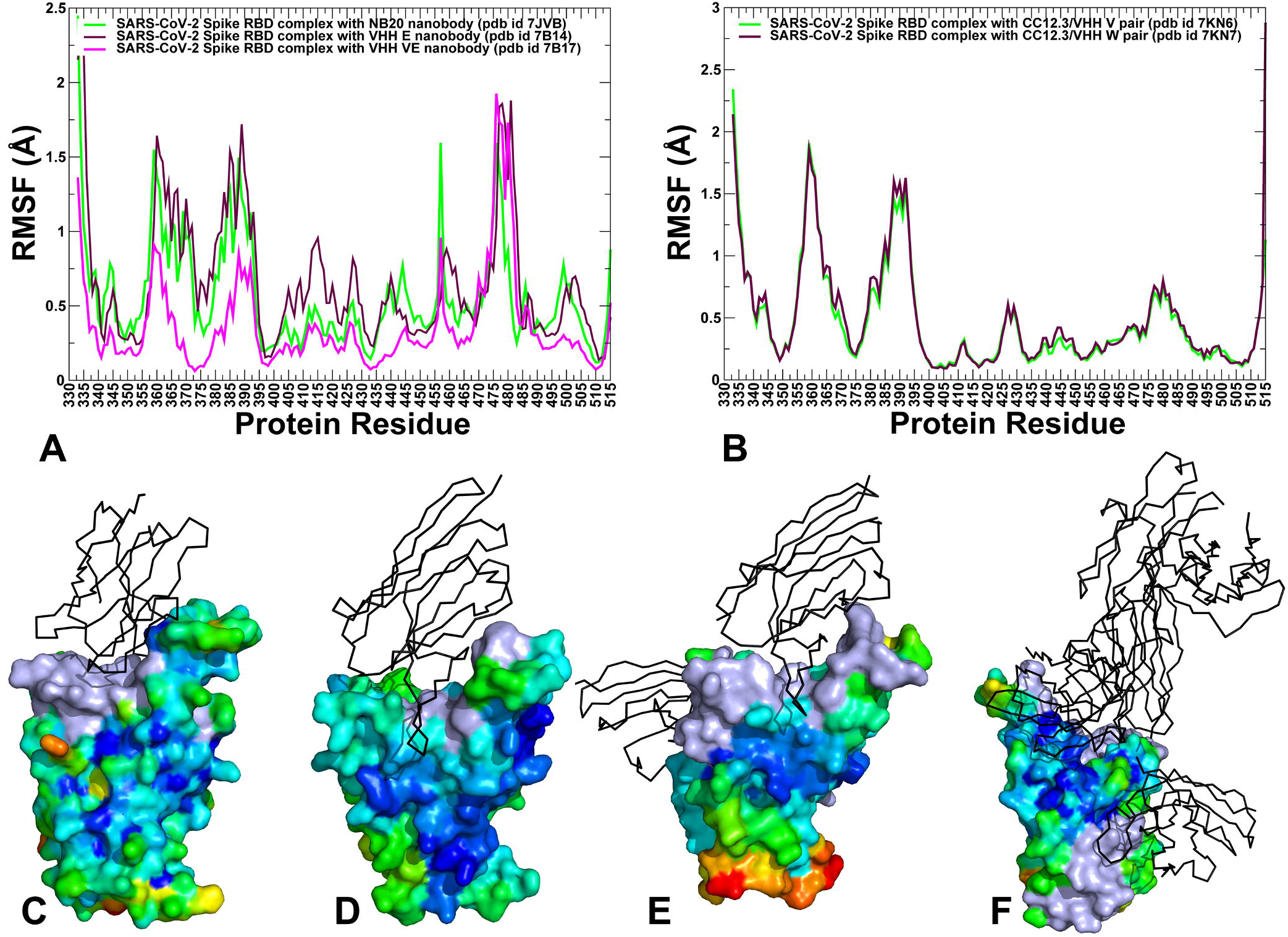
Conformational dynamics of the SARS-CoV-2 S-RBD. (A) The root mean square fluctuations (RMSF) profiles from MD simulations of the structures of the SARS-CoV-2 S-RBD complex with Nb20 (in green lines), VHH E (in maroon lines), and biparatopic nanobody VHH VE (in magenta lines). (B) The RMSF profile for the SARS-CoV-2 S-RBD complex with CC12.3 antibody and VHH V nanobody (in green lines), and combination of CC12.3 antibody and VHH W nanobody (in maroon lines). Structural maps of conformational mobility profiles for the S-RBD obtained from simulations of S-RBD complexes with Nb20 (C), VHH E (D), VHH VE (E), and CC12.3/VHH V (F). The conformational mobility level is shown from rigid (blue) to flexible (red). The bound nanobodies are shown in black-colored cartoons.

Interestingly, these regions include small α-helical segments (residues 367-370 and 383-388) that are tethered by C379-C432 and C391-C525 disulfide pairs. It is worth noting that the S-RBD region contains eight conserved cysteine residues, six of which form three disulfide linkages ( C336–C361, C379–C432 and C391–C525), which stabilize the β-sheet structure in the S-RBD protein. These α-helical segments are located close to the cryptic binding epitope (residues 369-384) targeted by VHH U/V/W nanobodies and appeared to undergo local functional movements that may help the S-RBD protein to accommodate nanobodies in the second epitope. In this context, the dynamics of the biparatopic nanobody complex VHH VE displayed the markedly reduced fluctuations in these intrinsically flexible regions that become stabilized and contribute to the binding interactions (Figure 2A). Another flexible RBD region that remained visibly mobile in the complexes is the tip of the RBM loop (residues 473-483) (Figure 2A). Interestingly, the immediately adjacent functional sites E484 and F486 also belong to this intrinsically mobile RBM region but are involved in binding contacts with nanobodies and become moderately stable in the complexes. In particular, Nb20 and VHH E binding can reduce mobility in the E484/F486 positions due to the favorable interactions (Figure 2A). Notably, these positions are commonly shared sites of nanobody-escaping mutations and only biparatopic nanobody construct becomes resilient to mutational escape in these positions. Another site of circulating variants N501 exhibited moderate fluctuations in the S-RBD complexes with Nb20 and VHH E, but was more stable in the complex with the biparatopic nanobody.

Hence, the conformational dynamics profiles highlighted two distal flexibility regions, one of which is associated with the RBM region and VHH E binding epitope, while another region overlaps with the cryptic RBD epitope targeted by VHH U/V/W nanobodies. According to our analysis, these flexible regions could play an important role in modulating S-RBD protein response to binding partners and mutational escape mechanisms. Importantly, the dynamics profiles revealed that the sites of circulating and common nanobody-escaping RBM mutations (L452, E484, F486, F490, and Q493) belong to the intrinsically adaptable flexible regions that become stable upon binding, particularly in the complex with the biparatopic nanobody.

Another important finding of the dynamics analysis is a significant stabilization of the S-RBD regions in the complex with a biparatopic construct VHH VE. Despite a similar interaction pattern of the RBM residues in the complex with the nanobody VHH E and a biparatopic analogue VHH VE, a significant fraction of the S-RBD (residues 400-465) become more stable in the complex with the biparatopic nanobody (Figure 2A). Structural map of the conformational mobility profile highlighted the broadly distributed immobilization of the S-RBD residues, particularly reflecting binding-induced constraints on the moving regions proximal to the binding epitopes, and leaving flexible only the peripheral RBD segment (residues 512-526) (Figure 2E). By rigidifying the moving RBD regions near the distal binding epitopes, the biparatopic nanobody VHH VE can suppress functional movements that are often exploited by escape variants to compromise the efficiency of binding interactions and limit neutralization potential. The simplest and most common way for viruses to escape nanobody blockage is by mutating residues in the easily accessible regions where viruses can tolerate mutations while escaping immune challenge. Our results suggest that the broad structural stability of the S-RBD regions induced by the biparatopic nanobody could counteract the mutational escape because simultaneous modifications in stable positions of two distinct and remote epitopes would tend to occur at a much lower rate than in a single RBM epitope.

We suggest that structural stability of the S-RBD induced by the VHH VE nanobody can also reflect avidity effects when multiple interactions in different epitopes can synergize to enhance the strength of protein-protein interactions in the multivalent complex, thus providing the thermodynamic advantage for stronger binding and neutralization capacity.^84, 85^ According to the experimental data, this nanobody demonstrated > 100-fold improved neutralizing activity and almost complete elimination of the escape mutants.^50^ Furthermore, avidity and positive allosteric cooperativity may work in concert^86^ when binding of a nanobody at the RBM epitope could increase affinity for a fused nanobody at the cryptic epitope. In the next chapters, using the ensemble-based mutational scanning of stability and binding, we will examine a hypothesis of potential synergies between binding of fused nanobodies that can be brought about via allosteric interactions and involve dynamic couplings between non-overlapping binding epitopes.

Conformational dynamics of the S-RBD complexes with CC12.3/VHH V and CC12.3/VHH W pairs showed a different pattern (Figure 2B). In some contrast, the entire RBD interface showed a considerable stabilization owing to a dense network of specific interactions formed by CC12.3 antibody. On the other hand, a very similar dynamics profile is seen in a second binding epitope targeted by VHH V and VHH W nanobodies (Figure 2B). Of particular importance, the stability of binding interactions formed by the loop in the distal end of RBD (residues 473-488). The antibody-induced stability of the distal loop is specific for CC12.3 binding as opposed to VHH E or Nb20 binding interactions. MD simulations revealed stable hydrogen network of interactions formed by pairs Y473 and S31, A475 and N32 and N487 and R94. Structural maps highlighted the greater rigidification of the RBM region in the complex with the CC12.3/VHH V combination pair, as compared to other complexes, indicating stronger modulation of stability by the antibody binding (Figure 2E, F).

Dynamic signatures of nanobody binding are also evident from the covariance matrixes of residue fluctuations suggesting conservation of correlated motions (Supporting Information, Figure S1). The correlation matrix of the S-RBD complex with VHH E showed that the S-RBD binding interface residues can be anti-correlated with fluctuations of VHH E residues. In the complex with the VHH VE biparatopic nanobody, the S-RBD dynamic couplings are generally stronger and positively correlated with fluctuations of the VHH V at the cryptic site, while mostly anti-correlated with VHH E (Supporting Information, Figure S1). The movements of VHH E and VHH V are dynamically strongly coupled and anti-correlated. Similarly, the positive correlations between fluctuations of the S-RBD and VHH V nanobody residues could be seen for the complex with CC12.3/VHH V complex, while weaker couplings were observed between S-RBD and CC12.3 motions (Supporting Information, Figure S1). Hence, strong positive correlations between S-RBD and VHH V residues supported the notion that binding to the second epitope could provide the dynamic “glue” for the biparatopic construct and strengthen the stability and regulatory role of the second binding interface as a global hinge of functional motions. At the same time, dynamic couplings between VHH V and VHH E may enable positive cooperativity in which VHH V binding arm incurs a small entropic loss and tight binding, allowing for the VHH E arm of the nanobody to maximize interactions with the RBM binding epitope.

### Nanobody-Induced Modulation of Functional Dynamics: Mutational Escape Sites Align with Hinge Centers and Cooperatively Moving Regions

We characterized collective motions for the SARS-CoV-2 S-RBD complexes averaged over low frequency modes using principal component analysis (PCA) of the MD trajectories (Figure 3). It is worth noting that the local minima along these profiles are typically aligned with the immobilized in global motions hinge centers, while the maxima correspond to the moving regions undergoing concerted movements leading to global changes in structure. The low-frequency ‘soft modes’ are often functionally important as mutations or binding partners tend to exploit and modulate the energetically favorable protein movements along the pre-existing slow modes to elicit specific protein response and induce allosteric transformations.

**Figure 3.**
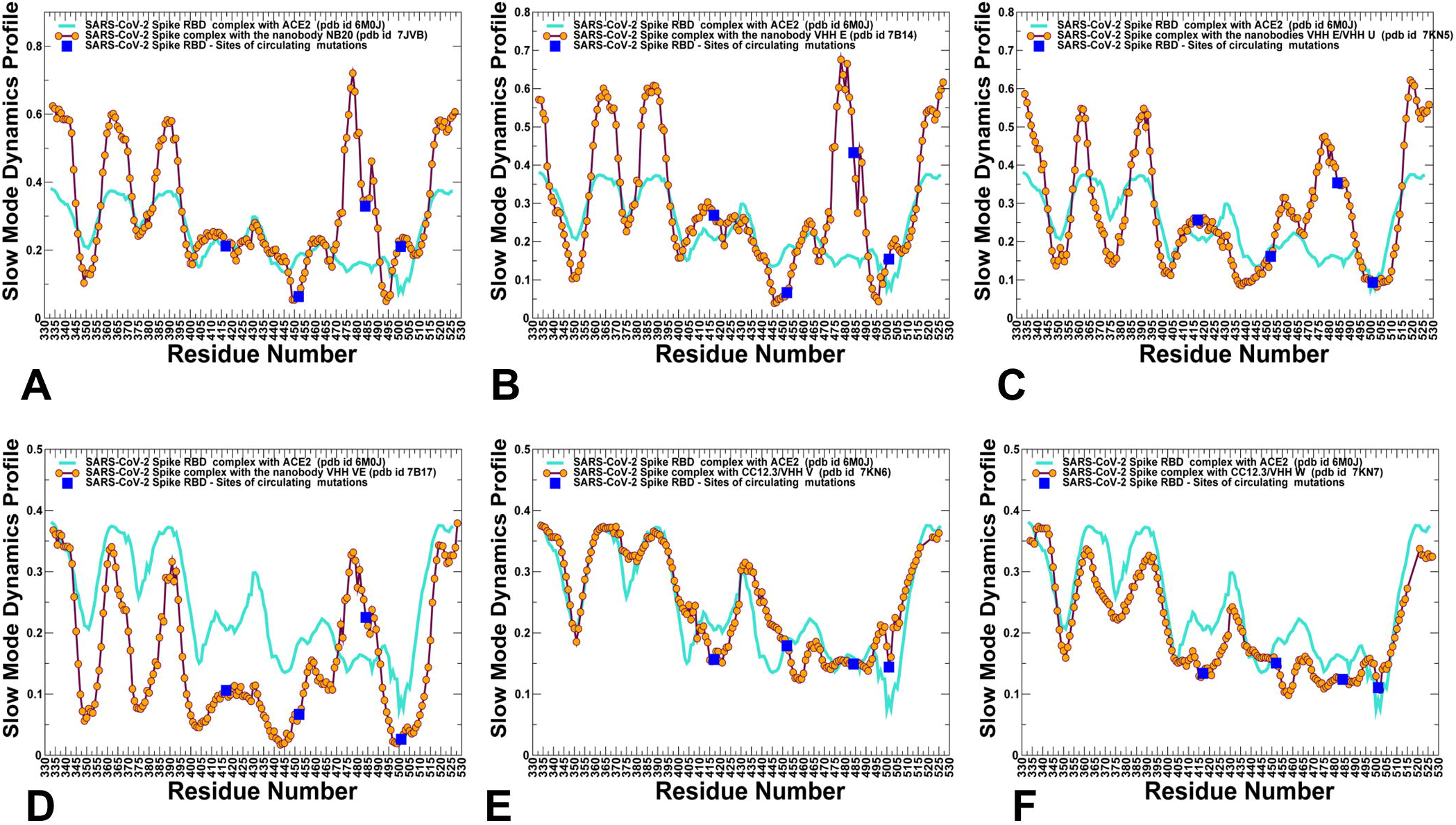
Collective dynamics profiles of the SARS-CoV-2 S complexes are shown for Nb20 nanobody, pdb id 7JVB (A), VHH E nanobody, pdb id 7B14 (B), VHH E/VHH U nanobody combination, pdb id 7KN5 (C), VHH VE biparatopic nanobody, pdb id 7B17 (D), CC12.3/VHH V antibody/nanobody combination, pdb id 7KN6 (E), and CC12.3/VHH W antibody/nanobody combination, pdb id 7KN7 (F). The profiles represent the mean square displacements in functional motions averaged over the three lowest frequency modes. The slow mode profiles are shown in maroon-colored lines with individual data points highlighted in filled orange circles. In each panel, the distributions are shown together with the slow mode profile for the S-RBD complex with ACE2 (in turquoise color lines).

The overall shape of the essential profiles was generally preserved between SARS-CoV-2 S complexes with nanobodies (Figure 3). We started by examining the S-RBD profiles for complexes with Nb20 nanobody (Figure 3A), VHH E nanobody (Figure 3B) and combinations of VHH E and VHH U nanobodies (Figure 3C). Nb20 and CC12.3 share class I epitopes that directly overlap with ACE2 and can potently inhibit the virus by competitively inhibit ACE2 binding, but also mimic ACE2 binding. The distributions revealed that sites of circulating variants K417 and N501 are aligned with local maxima corresponding to moving regions and L452 position belongs to the hinge region that is mostly immobilized in cooperative motions. Of particular notice is a common sharp bifurcated peak at the flexible region (residues 475-486) where large peak is aligned with 475-AGS-477 residues and another peak is at F486 position. At the same time, the periphery of the nanobody binding epitopes include E484 position that becomes partly stabilized and migrated in the distribution to a local minimum in this region.

Through this hinge-shift modulation nanobody binding may recruit E484 site into local hinge centers by reducing the flexibility of this residue. In the complex, E484 anchors cooperative movements of this dynamic region and therefore mutations in this site could affect the spectrum of functional nanobody motions. A comparison of S-RBD complexes with VHH E (Figure 3B), combination VHH VE/VHH U (Figure 3C) and biparatopic nanobody VHH VE (Figure 3D) pointed to several prominent well-defined hinge regions that are induced by binding. One of this hinge clusters is anchored by G447/Y449 residues and also includes L452 and L455 positions. This region corresponds to a sharp deep local minimum along the slow mode profile. Another noticeable hinge centers includes residues F490, L492, S494, G496, N501 and Y505. Both residue clusters are located in the relatively flexible regions of the S-RBD but become largely immobilized in the complexes. It could be noticed that the hinge clusters corresponded to pronounced and sharp local minima in the complex with the biparatopic nanobody (Figure 3D). These hinge clusters can mediate the motion in transferring perturbations throughout the system in a cascading fashion. Interestingly, the escape mutations in the VHH E interface featured G447S, Y449H/D/N, L452R, F490S, S494P/S, G496S, and Y508H.^50^ Hence, sites of escaping mutations at the VHH E interface could target the RBD positions that are involved in hinge clusters and therefore control collective motions in the complexes.

The residues in the dynamically moving region centered on the E484/F486 RBD positions are coupled to hinge sites and allow for movements to enable nanobody recognition. These functional sites corresponded to the local maxima of the essential mobility profiles in the complexes with VHH E/VHH U (Figure 3C) and VHH VE biparatopic nanobody (Figure 3D). As a result, this flexible region can experience functional movements and contribute to allosteric conformational changes induced by binding.

An important finding of this analysis is that escape mutations at the RBM epitope targeted by VHH E may arise in the intrinsically dynamic positions that play an important regulatory role in coordination and execution of collective functional motions along pre-existing slow modes. We argue that these mutations may change the distribution of regulatory centers and alter global functional movements of nanobodies required for binding and activity. We also observed that structurally rigid residues in the cryptic binding epitope including Y369, S371, F374, and F377 are aligned with conserved hinge positions. The hinge clusters formed at the two distant binding epitopes may work cooperatively to regulate functional dynamics of the biparatopic nanobody. Combined with the conformational dynamics analysis, the results suggested that allosteric couplings between these regulatory clusters could collectively determine functional movements and mechanism of the nanobody binding.

The slow mode profiles for the S-RBD complexes with CC12.3/VHH V and CC12.3/VHH W pairs showed a distinct pattern (Figure 3E, F). We observed a very shallow minima region in the RBM binding epitope where E484 and N501 positions are aligned with largely immobilized in collective movements regions. This analysis also indicated that the experimentally known sites of escaping mutations Y421, L455, F456 and Y508 for CC12.3 antibody^49^ featured among shallow minima of the profile. In this model, the emergence of broad and shallow local minima regions may signal tolerance to mutations at individual positions as these changes would not significantly affect the pattern of functional dynamics as opposed to profiles featuring several sharp hinges that are more vulnerable to mutations.

### Deep Mutational Scanning Identifies Structural Stability and Binding Affinity Hotspots in the SARS-CoV-2 Complexes and Explains Patterns of Nanobody-Escaping Mutations

Using the ensemble-based mutational scanning of binding, we systematically modified the S-RBD residues and computed the corresponding binding free energy changes for studied SARS-CoV-2 S-RBD complexes with nanobodies (Figures 4, 5). The resulting mutational heatmaps provided a convenient visual representation of in silico screening, pointing to the S-RBD residues with a high sensitivity to modifications. In our model, these sites are considered as potential binding energy hotspots. Concurrently, we also employed the FoldX approach with the all-atom representation of protein structure^87–90^ to evaluate the folding free energies for the S-RBD residues in the complexes and identify hotspots of protein stability. The protein stability ΔΔG changes were computed by averaging the results of computations over 1,000 samples obtained from MD simulation trajectories.^91, 92^ In this analysis, positive folding free energy contributions are suggestive of stability weaknesses sites and negative folding free energies point to the structurally stable positions in the S-RBD.

**Figure 4.**
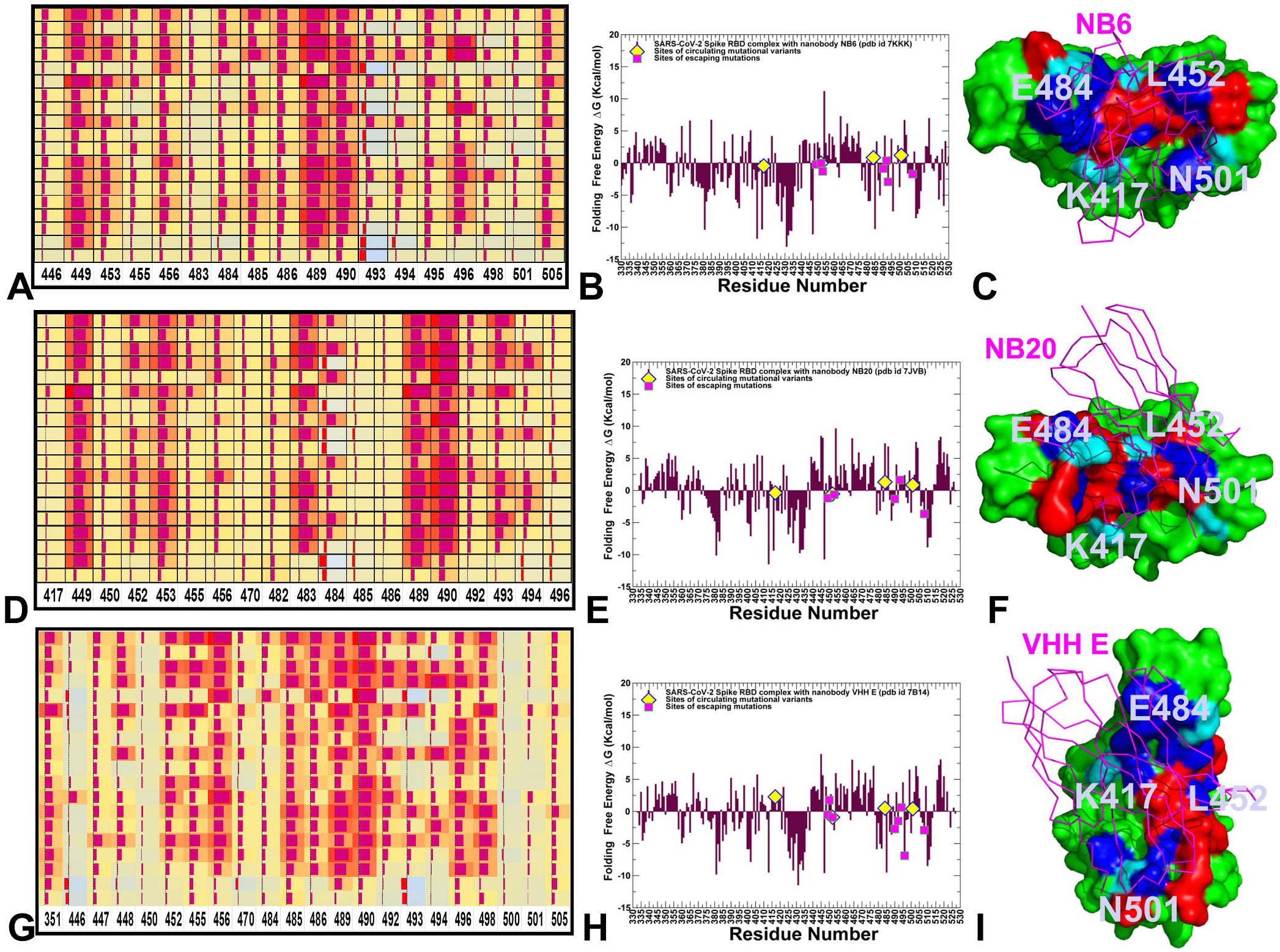
The mutational scanning heatmap for the SARS-CoV-2 S complex with Nb6 nanobody, pdb id 7KKK (A). The residue-based folding stability analysis of the SARS-CoV-2 S-RBD complex with Nb6 (B). Structural map of the S-RBD binding epitope in the complex with Nb6 (C). The binding energy hotspots correspond to residues with high mutational sensitivity. The S-RBD is shown in green surface. The bound nanobody is shown in magenta-colored cartoons. The epitope residues are shown in red and the binding energy hotspots from are shown in blue surface. The positions of K417, L452, E484 and N501 are shown in light cyan color and annotated. The mutational scanning heatmap for the SARS-CoV-2 S complex with Nb20 nanobody, pdb id 7JVB (D). The residue-based folding stability analysis of the S-RBD complex with Nb20 (E). Structural map of the S-RBD binding epitope in the complex with Nb20 (F). The mutational scanning heatmap for the SARS-CoV-2 S complex with VHH E nanobody, pdb id 7B14 (G). The residue-based folding stability analysis of the S-RBD complex with VHH E nanobody (H). Structural map of the S-RBD binding epitope in the complex with VHH E nanobody (I). The S-RBD is shown in green surface. The bound nanobody is shown in magenta-colored cartoons. The epitope residues are shown in red and the binding energy hotspots are shown in blue surface. The positions of K417, L452, E484 and N501 are shown in light cyan color and annotated. The heatmaps show the computed binding free energy changes for 19 single mutations on the binding epitope sites. The squares on the heatmap are colored using a 3-colored scale - from light blue to red, with red indicating the largest destabilization effect. The data bars correspond to the binding free energy changes, with positive destabilizing value shown by bars towards the right end of the cell and negative stabilizing values as bars oriented towards left end.

**Figure 5.**
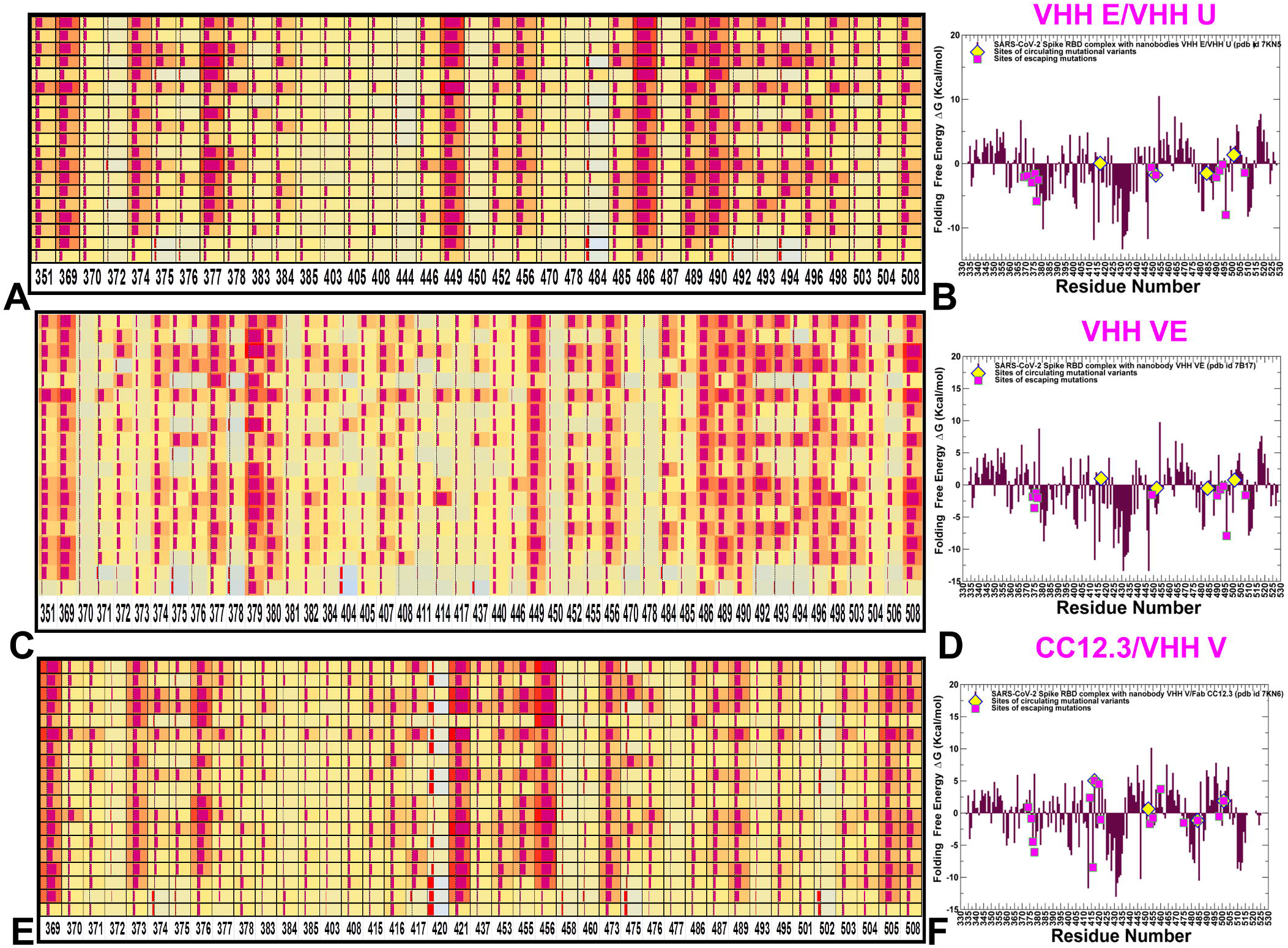
The mutational scanning heatmap for the SARS-CoV-2 S complex with VHHE/VHH U combination of nanobodies, pdb id 7KN5 (A). The residue-based folding stability analysis of the SARS-CoV-2 S-RBD complex with VHH E/VHH U combination (B). The binding energy hotspots correspond to residues with high mutational sensitivity. The mutational scanning heatmap for the SARS-CoV-2 S complex with VHH VE biparatopic nanobody, pdb id 7B17 (C). The residue-based folding stability analysis of the S-RBD complex with VHH VE (D). The mutational scanning heatmap for the SARS-CoV-2 S complex with CC12.3/VHH V antibody/nanobody combination, pdb id 7KN6 (E). The residue-based folding stability analysis of the S-RBD complex with CC12.3/VHH V antibody/nanobody combination (F). The heatmaps show the computed binding free energy changes for 19 single mutations on the binding epitope sites. The squares on the heatmap are colored using a 3-colored scale - from light blue to red, with red indicating the largest destabilization effect. The data bars correspond to the binding free energy changes, with positive destabilizing value shown by bars towards the right end of the cell and negative stabilizing values as bars oriented towards left end.

We first analyzed the mutational profiles for the S-RBD complexes with Nb6 and Nb20 nanobodies (Figure 4). Mutational sensitivity analysis of the S-RBD binding with Nb6 showed that a significant fraction of the epitope residues can contribute to the binding affinity, suggesting that the nanobody interactions with the ACE2-binding site may be highly efficient (Figure 4A). The key binding energy hotspots for Nb6 corresponded to the hydrophobic residues Y449, L453, L455, F456, G485, Y489, F490, G496 and Y505 (Figure 4A,B). A number of these positions are recognized as important binding affinity hotspots for ACE2 as evident from deep mutagenesis scanning of SARS-CoV-2 interactions with the ACE2 host receptor.^39^ The interaction pattern and similarity in the binding energy hotspots with ACE2 supported the notion of structural mimicry that may be efficiently exploited by Nb6 nanobody to competitively inhibit the ACE binding region.

The folding free energies of the S-RBD residues reflected high stability of the conserved core of the S-RBD consisting of antiparallel β strands (β1 to β4 and β7) (residues 354-358, 376-380, 394-403, 431-438, 507-516) and also pointed to a moderate stabilization of β5 and β6 (residues 451-454 and 492-495) that link the binding region to the central core (Figure 4B). Of interest are negative folding free energies for a range of binding energy hotspots Y449, L455, F456, G485, Y489, and Y505 indicating that these positions correspond to the conserved stability centers. Accordingly, escape mutations are unlikely to target these structurally stable positions as this would adversely affect not only nanobody binding but also the RBD folding stability that is required for binding to the host receptor. Structural map of the binding epitope residues and binding energy centers showed that sites of circulating variants K417, L452, E484 and N501 belong to the edges of the RBM binding epitope and are located in the immediate proximity of the hotspot residues (Figure 4C). The mutational sensitivity map also shed some light on structure-functional role of sites targeted by common resistant mutations (F490S, E484K, Q493K, F490L, F486S, F486L, and Y508H) that evade many individual nanobodies.^47^ Indeed, we found that E484, F486, and F490 positions can be sensitive to Nb6 binding (Figure 4A-C), while variations in these sites are known to have only minor effect on ACE2 binding.

For the S-RBD complex with Nb20 nanobody, the binding energy hotspots corresponded to residues Y449, Y453, V483, Y489, F490, and Q493 (Figure 4D). A several other weaker hotspot centers are associated with L452, E484, L492 and Y494 positions. Despite some differences in the binding epitope composition, there are a number of commonly shared hotspots for Nb6 and Nb20 nanobodies including Y449, F456, Y489, and F490 (Figure 4D). At the same time, some Nb20-specific binding hotspots corresponded to V483; L492 and Q493 positions (Figure 4D). Interestingly, mutations of E484 residue (E484K/D/N) could also result in a significant loss of binding affinity (∼2.0 kcal/moll) for both Nb6 and Nb20 nanobodies, while the respective modifications lead to the improved binding with the host receptor (Supporting Information, Figure S2). The emergence of Q493 hotspot is due to consistent hydrogen bonding with A29 and R97 of Nb20 observed in the equilibrium ensemble, while F490 can establish similarly sustainable interactions with R31 of Nb20 nanobody. These findings are consistent with the original structural studies showing E484 on the RBD is the “Achilles heel” of the ultra-potent Nb20 and Nb21 that can dramatically reduce the ultrahigh affinity of these class I nanobodies.^45, 46^ In simulations, Nb20 maintains hydrogen bonding and stabilizing electrostatic interactions with E484 using side chains of R31, R97 and Y104 residues, so that circulating variant mutation E484K can alter the pattern of these contacts and make them repulsive, leading to the reduced binding affinity. E484 also mediates a network with neighboring RBM residues such as F490, F489, N487, Y486, and V483 to participate in binding. The E484K mutation can substantially destabilize the interface packing by electrostatic repulsion with R31, subsequently disrupting the cation-π stacking interaction between R31 and F490 residues (Supporting Information, Figure S2). Consistent with the results of mutational scanning, the engineered charge reversal on R31 (R31D) is unable to recover the salt bridge and restore binding to the E484K mutant.^45, 46^

Notably, the sites of nanobody-escaping mutations featured a relatively moderate stability as evident from small negative folding free energies (Figure 4E, F). The sites of circulating mutations K417, L452, E484 and N501 displayed small positive values, indicating that these positions correspond to weak points of protein stability. The marginal stability for sites of escaping and circulating mutations is consistent with the notion that virus tends to target positions where mutations would not appreciably perturb the RBD folding stability that is prerequisite for proper activity of spike protein and binding with the host receptor. By targeting dynamic and structurally adaptable hotspots such as E484, F486, and F490 that are relatively tolerant to mutational changes, virus tends to exploit conformational plasticity in these regions in eliciting specific escape patterns that would impair nanobody binding.

To further test this conjecture, we examined the binding free energy changes associated with common variants N501Y, K417N/T, E484K and L452R (Supporting Information, Figure S2). In the complex with Nb20, E484D/K/Q mutations appeared to be highly destabilizing resulting respectively in ∼2.77, 2.83 and 2.63 kcal/mol losses in the binding affinity (Supporting Information, Figure S2). At the same time, N501Y mutation led to the moderately increased binding affinity which is consistent with the respective improvements in affinity to ACE2. In addition, known escaping mutants F490S, F490L, Q493K resulted in large destabilization changes (>2.5 kcal/mol) which is in agreement with deep mutagenesis results showing that these variants likely to cause immune escape.^45^ The data also revealed moderate sensitivity of Nb20 binding to mutations in L452 site, where circulating variants L452R and L452K triggered an appreciable loss of affinity (∼1.0 kcal/mol). Consistent with the experimental studies, mutational scanning analysis demonstrated that F490, E484, F486, and Q493 positions are sites of escaping mutations Nb6 and Nb20 nanobodies.

Structural mapping of protein stability hotspots for Nb6 and Nb20 nanobodies highlighted two distinct conserved clusters of residues localized at the center of the RBM binding epitope and in the S-RBD core region near the cryptic binding epitope (Supporting Information, Figure S3A,B). This structural distribution of the energetic hotspots is preserved in the complexes with nanobodies, indicating that protein stability centers are mainly localized near the cryptic binding epitope exploited by nanobody cocktails. The energetic hotspots at the second binding epitope correspond to conserved rigid residues that are unlikely to emerge as mutational escape positions due to functional requirements for RBD folding. At the same time, the RBM hotspots are localized near functionally adaptable regions and emerge due to nanobody-induced stabilization and binding affinity (Supporting Information, Figure S3).

For the S-RBD complex with VHH E nanobody targeting the RBD region, a consistent binding footprint revealed several clusters of binding energy hotspots (Figure 4G-I). VHH E shared a number of common hotspot centers with Nb6 and Nb20 nanobodies, reflecting structural and energetic importance of key binding energy hotspots in the RBD region (Figure 4G). One group of these energetic centers corresponded to sites L452, Y453 and F456, while the second cluster included functional positions G485, F486, Y489, F490, L492, G496 and Q498. Strikingly, a comparison of the binding epitopes and hotspots for RBB-targeting nanobodies revealed that sites of common escaping mutations L452, E484, F486 and F490 are characterized by high mutational sensitivity (Figure 4G-I). It should be noted that sites of circulating variants are characterized by only marginal stability or weakly destabilizing, while residues targeted by escaping mutations at VHH E epitope can be moderately stable (Figure 4H). A the same time, mutations in E484 functional site produced considerable destabilization changes in the binding affinity for VHH E complex with the S-RBD (Supporting Information, Figure S4), indicating that this critical position can be sensitive to variations that compromise the nanobody binding interactions.

Overall, this analysis indicated that binding energy hotspots in the nanobody complexes with Nb6, Nb20 and VHH E are determined by balance between protein stability constraints and binding interactions with the RBM binding epitope. Importantly, the nanobody-escaping RBD mutations typically produce moderate but still appreciable changes in the protein stability, while global circulating variants appeared to have a minor effect on the RBD stability, allowing virus to evade nanobody binding without compromising folding constraints and fitness for the host receptor.

### Binding Footprints for Nanobody Combinations at Distinct Epitopes Reveal Changes in the Mutational Landscape and Increased Resilience to Escape Mutations

We also identified binding affinity fingerprints for the SARS-CoV-2 S-RBD complexes with a nanobody combination VHH E/VHH U targeting different epitopes (Figure 5A,B), and biparatopic nanobody VHH VE that has two antigen-binding sites in one molecule and obtained by fusing nanobodies that targeted distinct epitope regions (Figure 5C,D). Through ensemble-based deep mutational profiling of stability and binding affinities, we identify critical hotspots and examine molecular mechanisms of mutational escape. In this analysis, we test the hypothesis that through cooperative dynamic and energetic changes, nanobody combinations and biparatopic nanobody can elicit synchronized changes in the binding interactions and induce the increased resilience to common escape mutants.

For S-RBD complex with the nanobody pair VHHE/VHH U targeting two different epitopes, the panel of RBD residues involved in the favorable binding interactions increased, but the number of binding energy hotspots remained similar to the complexes with single nanobodies (Figure 5A). The S-RBD hotspot residues were aligned with the same group of residues Y449, F486, Y489, F490, Y508 (Figure 5A). In agreement with the experiments^50^, mutations at the VHH E interface Y449H/D/N, F490S, S494P/S, G496S, and Y508H produced destabilizing ΔΔG changes exceeding 2.0 kcal/mol (Figure 5A,B). A number of hotspot positions were also observed in the second cryptic epitope including conserved and stable residues Y369, S371, F374 and F377. Our mutational scanning analysis also showed that escaping mutations Y369H, S371P, F374I/V, T376I, F377L, and K378Q/N at the VHH U interface resulted in considerable destabilization losses with ΔΔG ∼2.5 kcal/mol/ (Figure 5A,B).

In the complex with the biparatopic nanobody VHH VE spanning two binding epitopes (Figure 5C,D), the pattern remained generally similar, highlighting a strong contribution of conserved stability centers F377, C379 and Y489 as well as RBD hotspots Y449, F486, L490 and L492. A more detailed analysis of the binding energetics provided several important quantitative insights. First, we noticed a significant number of the binding epitope residues interacting with VHH V segment of the nanobody (Figure 5C). The binding interactions in this epitope are determined by RBD core residues 369-384. Of notice are strong contributions of F377, C379, Y380 residues acting as rigid binding energy hotspots (Figure 5C). Second, for the S-RBD complexes with VH E/VHH U combination and VHH VE biparatopic nanobody many of the hotspot centers corresponded to conserved centers of protein stability where mutations could affect the RBD folding and therefore unlikely to emerge as nanobody-escaping variants.

Even though these nanobodies make contacts with sites of circulating variants K417, L452 and E484, our results showed that these positions can be quite tolerant to mutations (Supporting Information, Figure S4). Interestingly, mutations L452R and T478K of B.617.2 lineage that can induce escape to neutralizing antibodies targeting the NTD and RBD^63, 64^ resulted in relatively moderate binding energy losses for the biparatopic nanobody (Figure 5C,D). This is consistent with the observed effectiveness of the nanobodies against these antigenic variants.^50^ The results also indicated that only moderately flexible RBD sites F486 and F490 are consistently featured as common binding energy hotspots for these complexes which may explain why escape mutants in these positions are known to dominate at the VHH E interface.^50^

The important finding of this analysis is that mutational sensitivity of the binding hotspots in the RBM epitope can be attenuated in the complex with the biparatopic nanobody (Figure 5C). This may contribute to the increased resilience of the VHH VE engineered nanobody to mutational escape. These results reflect dynamic intermolecular couplings of the S-RBD with the VHH V arm that can effectively anchor the biparatopic nanobody at the cryptic epitope providing thermodynamic advantage for binding of VHH E at the RBM epitope. Structural map of the binding epitopes in the S-RBD complexes with VHH E/VHH U combination and VHH VE biparatopic nanobody highlighted preservation of the key binding energy hotspots in the RBM epitope (Figure 6). We also observed that sites of circulating variants K417, L452, E484 and N501 are localized close to the hotspot residues and could form structural “borders” of the binding epitopes (Figure 6). This visual characterization of the RBM binding footprints suggested that small variations in the sites of circulating variants would have a moderate effect on binding provided that corresponding structural perturbations induced by mutations are local and could be tolerated. A comprehensive structural mapping of the energy hotspots in the complexes with the nanobodies revealed a large and dense cluster of evolutionary conserved stability centers at the second binding epitope (Supporting Information, Figure S3).

**Figure 6.**
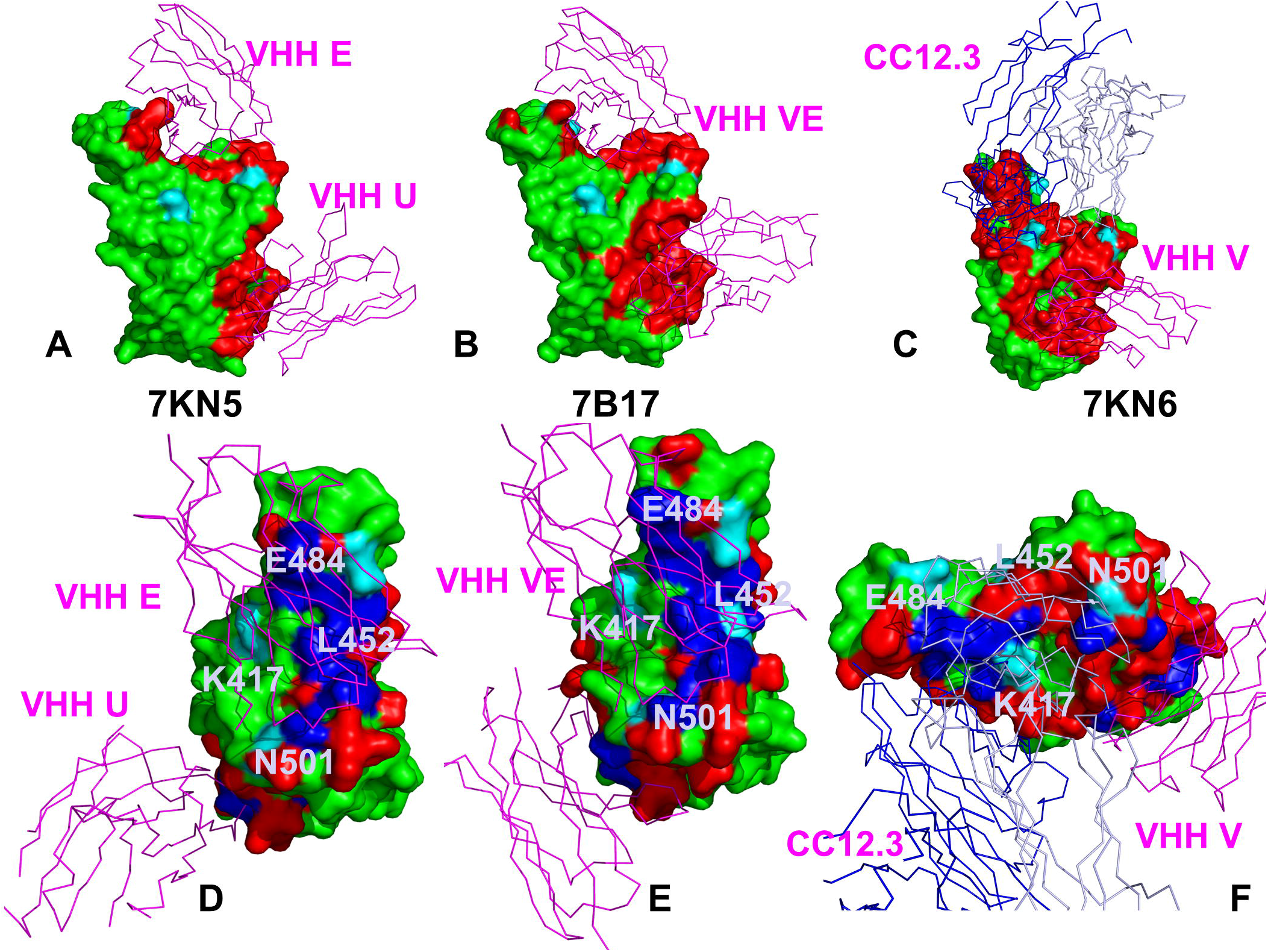
The structure of the S-RBD complex with VHH E/VHH U combination, pdb id 7KN5 (A), complex with biparatopic nanobody VHH VE (B), and CC12.3/VHH V antibody/nanobody combination (C). S-RBD is shown in green surface. The epitope residues are shown in red. The positions of K417, L452, E484 and N501 are shown in light cyan color. The bound nanobodies are shown in magenta-colored cartoons. Top views of the binding epitopes for S-RBD complexes with VHH E/VHH U combination (D), complex with biparatopic nanobody VHH VE (E), and CC12.3/VHH V antibody/nanobody combination (F). The S-RBD is shown in green surface. The epitope residues are shown in red and the binding energy hotspots are shown in blue surface. The positions of K417, L452, E484 and N501 are shown in light cyan color and annotated.

This stable anchor could be used by the biparatopic nanobody as a regulatory anchor that firmly immobilizes VHH U and VHH V nanobodies at the cryptic binding site and exerts allosteric control over structural changes in the RBM epitope. Accordingly, due to avidity effects, binding of the VHH E arm at the RBM epitope may incur a smaller entropic cost and allow for local structural accommodations in these regions to compensate for the loss of binding interactions with the mutants. This suggests a plausible mechanism by which biparatopic nanobodies can leverage dynamic couplings mediated by the rigid nanobody arm to synergistically inhibit distinct binding epitopes and suppress mutational escape.

The mutational heatmaps of the S-RBD binding with antibody CC12.3 combinations with VHH V and VHH W nanobodies (Figure 5E,F) recapitulated the experimentally known sites of escaping mutations Y421, L455, F456 and Y508 for CC12.3.^49^ The cryptic binding epitope featured a number of rigid binding hotspots in the RBD core corresponding to residues 369-368. In the RBM epitope, the binding hotspots were broadly distributed and corresponded to Y421, Y453, F456, Y473, Y489, and Y505 sites (Figure 5E). These sites also serve as conserved structural stability centers (Figure 5F) and are unlikely to be selected for mutational escape due to their role in the S-RBD protein folding and stability. Importantly, this analysis showed that binding of CC12.3/VHH V(W) combinations can tolerate mutations in sites of circulating variants K417, L452, E484, and N501, yielding only small energy differences (Supporting Information, Figure S4), which is consistent with the observed resilience of these cocktails against these antigenic variants. The results confirmed that nanobody combinations could limit the emergence of escape mutants that confer resistance to specific regions centered on F456, F490 and Q493 residues.

We also hypothesized that mechanisms of structural mimicry may employed by the nanobodies to compete with ACE2 binding. To test this conjecture, we evaluated the correlations between binding free energy changes of the S-RBD residues with ACE2 and studied nanobodies (Figure 7). A consistent significant correlation (Pearson correlation coefficient R ∼ 0.7-0.85) was found between the effects of RBD mutations on binding with ACE2 and the nanobodies. This indicated that the nanobodies can efficiently mimic the binding energetics of the host receptor. We specifically highlighted the binding free energy changes associated with mutations at sites of circulating variants K417, L452, E484, and N501 (Figure 7). Interestingly, while mutations in these positions tend to cause similar and moderate binding energy changes for ACE2 and nanobodies, we noticed some dispersion of the energetic changes showing a larger destabilization effect on nanobody binding (∼ 1.0 −1.5 kcal/mol) than on ACE2 binding (∼ 0.5 kcal/mol). This effect was similar for the VHH E nanobody interacting at the RBM epitope (Figure 7A), the nanobody combination (Figure 7B), and biparatopic nanobody VHH VE (Figure 7C). Hence, modifications in these positions incur moderate binding losses at the VHH E binding site that could prevent complete mutational escape. This analysis revealed a very strong correlation between binding free energy changes in these functional positions with ACE2 and CC12.3/VHH V combination (Figure 7D). In this case, we found a high degree of tolerance to mutations in sites of circulating variants, suggesting that a combination of antibody and nanobody binding may be very efficient in a broad suppression of mutational escape. Structural and dynamic mimicry of the protein stability and binding interactions by the nanobody combinations could limit the escaping potential for the virus in sites of common circulating variants.

**Figure 7.**
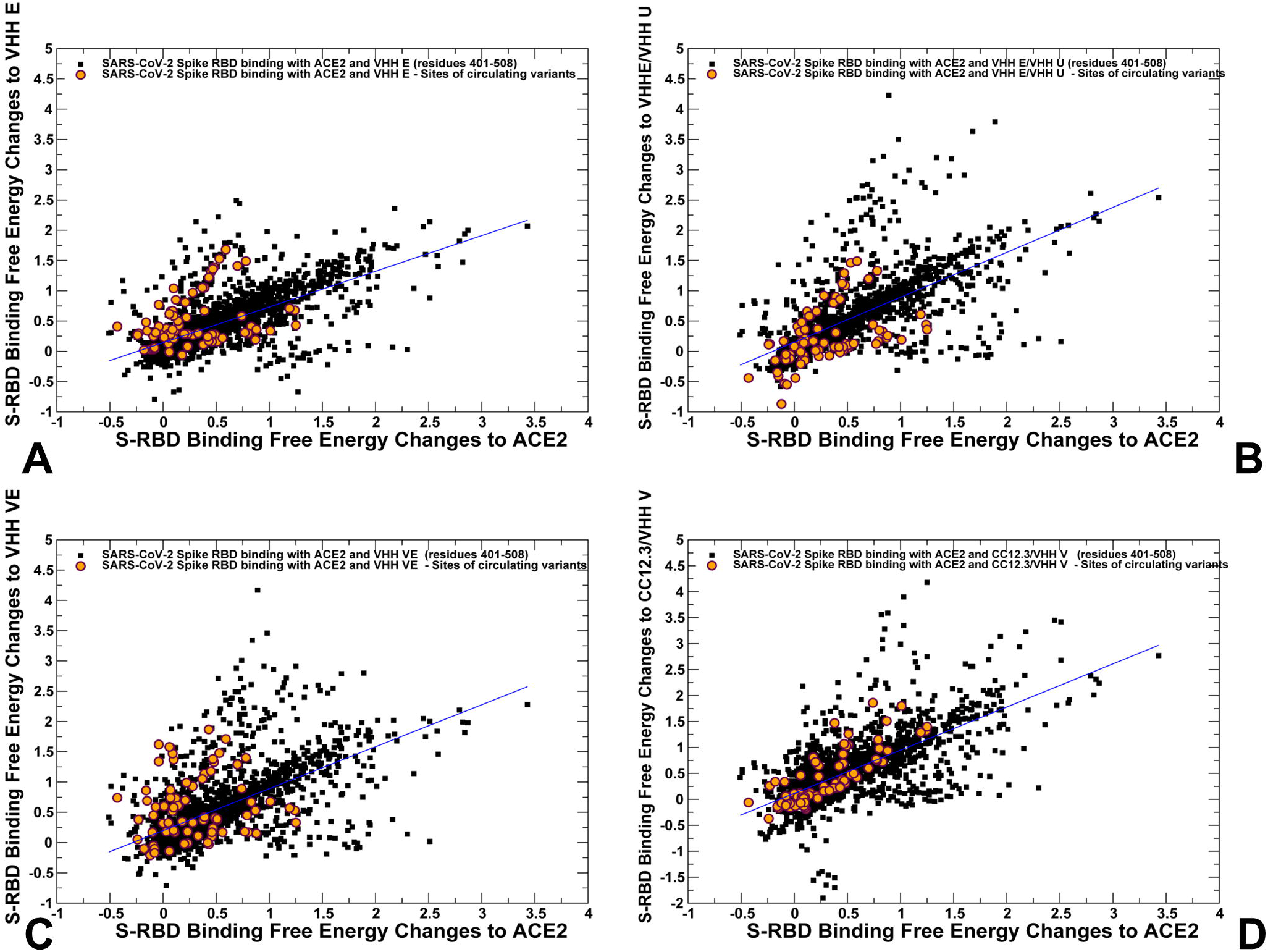
The 2D scatter plots of binding free energy changes for the S-RBD residues in complexes with ACE2 and nanobodies. (A) The scatter plot of binding free energy changes from deep mutational scanning of the S-RBD residues with ACE2 and VHH E nanobody. (B) The scatter plot of binding free energy changes from deep mutational scanning of the S-RBD residues with ACE2 and VHH E/VHH U combination. (C) The scatter plot of binding free energy changes from deep mutational scanning of the S-RBD residues with ACE2 and biparatopic nanobody VHH VE. (D) The scatter plot of binding free energy changes from deep mutational scanning of the S-RBD residues with ACE2 and CC12.3/VHH V combination. The data points are shown as black-filled squares and scatter points corresponding to the binding free energy changes in sites K417, L452, E484 and N501 are shown as orange-filled circles.

## Conclusions

We combined atomistic simulations with the ensemble-based mutational profiling of binding interactions for a diverse panel of SARS-CoV-2 S-RBD complexes with nanobodies. Conformational dynamics profiles revealed modulation of structural stability in the S-RBD regions induced by the nanobody combinations and biparatopic nanobody, particularly a significant rigidification of the binding interface a the cryptic binding epitope. The analysis of collective dynamics in the nanobody complexes showed that escape mutations at the RBM epitope may emerge in the intrinsically dynamic positions that become stabilized upon binding and play an important regulatory role in coordination of functional motions. Using ensemble-based deep mutational profiling of stability and binding affinities, we identified critical hotspots and characterized molecular mechanisms of SARS-CoV-2 S binding with single nanobodies, nanobody cocktails and biparatopic nanobodies. Deep mutational scanning showed that the pattern of escaping mutations to nanobody binding may be restricted to several common dynamic sites. The results also revealed that the biparatopic nanobody can use binding at the cryptic binding site to establish a structurally stable regulatory “anchor” that mediates dynamic couplings between nanobody arms and provides allosteric control over structural changes in the RBM epitope. This suggests a plausible mechanism by which biparatopic nanobodies can leverage dynamic couplings to synergistically inhibit distinct binding epitopes and suppress mutational escape. We argue that mutational escape mechanisms may be generally controlled and manipulated through structurally and energetically adaptable binding hotspots in the RBM epitope that are dynamically coupled to stability centers in the cryptic epitope.

## Methods

### Structure Preparation and Analysis

The cryo-EM structures of SARS-CoV-2 S proteins used in this study included the S-RBD complex with ACE2 (pdb id 6M0J), and crystal structures of the SARS-CoV-2 S-RBD complexes with a series of highly potent neutralizing nanobodies and various nanobody cocktails including Nb6 (pdb id 7KKKJ), Nb20 (pdb 7JVB), VHH E (pdb id 7B14), a nanobody combination pair VHH E/ VHH U (pdb id 7KN5), biparatopic nanobody VHH VE (pdb id 7B17), as well as antibody/nanobody cocktails such as combination of CC12.3 antibody and VHH V nanobody (pdb id 7KN6), and combination of CC12.3 antibody and VHH W nanobody (pdb id 7KN7) (Figure 1). All structures were obtained from the Protein Data Bank.^93, 94^ Hydrogen atoms and missing residues were initially added and assigned according to the WHATIF program web interface.^95^ The structures were further pre-processed through the Protein Preparation Wizard (Schrödinger, LLC, New York, NY) and included the check of bond order, assignment and adjustment of ionization states, formation of disulphide bonds, removal of crystallographic water molecules and co-factors, capping of the termini, assignment of partial charges, and addition of possible missing atoms and side chains that were not assigned in the initial processing with the WHATIF program. The missing loops in the cryo-EM structures were also reconstructed using template-based loop prediction approaches ArchPRED.^96^ The side chain rotamers were refined and optimized by SCWRL4 tool.^97^ The protein structures were subsequently optimized using atomic-level energy minimization with a composite physics and knowledge-based force fields using 3Drefine method.^98^

The shielding of the receptor binding sites by glycans is an important feature of viral glycoproteins, and glycosylation on SARS-CoV-2 proteins can camouflage immunogenic protein epitopes.^99, 100^ While all-atom MD simulations with the explicit inclusion of the complete glycosylation shield provide a rigorous and the most detailed description of the conformational landscape for the SARS-CoV-2 S proteins, such simulations remain to be computationally highly challenging due to the size of a complete SARS-CoV-2 S system. The structure of glycans at important glycosites (N122, N165, N234, N282, N331, N343, N603, N616, N657, N709, N717, N801, N1074, N1098, N1134, N1158) of the closed and open states of SARS-CoV-2 S protein was previously determined in the cryo-EM structures of the SARS-CoV-2 spike S trimer in the closed state (pdb id 6VXX) and open state (pdb id 6VYB). These glycans were incorporated in simulations of the SARS-CoV-2 S protein complexes in addition to the experimentally resolved glycan residues present in the cryo-EM structures of studied SARS-CoV-2 S proteins.

### MD Simulations

All-atom 1µs MD simulations have been performed for all studied protein structures. The structures of the SARS-CoV-2 S trimers were simulated in a box size of 85 Å × 85 Å × 85 Å with buffering distance of 12 Å. Assuming normal charge states of ionizable groups corresponding to pH = 7, sodium (Na+) and chloride (Cl-) counter-ions were added to achieve charge neutrality and a salt concentration of 0.15 M NaCl was maintained. All Na^+^ and Cl^-^ ions were placed at least 8 Å away from any protein atoms and from each other. All-atom MD simulations were performed for an N, P, T ensemble in explicit solvent using NAMD 2.13 package^101^ with CHARMM36 force field.^102^ Long-range non-bonded van der Waals interactions were computed using an atom-based cutoff of 12 Å with switching van der Waals potential beginning at 10 Å. Long-range electrostatic interactions were calculated using the particle mesh Ewald method^103^ with a real space cut-off of 1.0 nm and a fourth order (cubic) interpolation. Simulations were run using a leap-frog integrator with a 2 fs integration time step. All atoms of the complex were first restrained at their crystal structure positions with a force constant of 10 Kcal mol^-1^ Å^-2^. Equilibration was done in steps by gradually increasing the system temperature in steps of 20K starting from 10K until 310 K and at each step 1ns equilibration was done keeping a restraint of 10 Kcal mol-1 Å-2 on the protein C_α_ atoms. After the restrains on the protein atoms were removed, the system was equilibrated for additional 10 ns. An NPT production simulation was run on the equilibrated structures for 500 ns keeping the temperature at 310 K and constant pressure (1 atm). In simulations, the Nose–Hoover thermostat^104^ and isotropic Martyna–Tobias–Klein barostat^105^ were used to maintain the temperature at 310 K and pressure at 1 atm respectively.

We performed principal component analysis (PCA) of reconstructed trajectories derived from CABS-CG simulations using the CARMA package^106^ and also determined the essential slow mode profiles using elastic network models (ENM) analysis.^107–109^ Two elastic network models: Gaussian network model (GNM) and Anisotropic network model (ANM) approaches were used to compute the amplitudes of isotropic thermal motions and directionality of anisotropic motions. The functional dynamics analysis was conducted using the GNM in which protein structure is reduced to a network of *N* residue nodes identified by *C_α_* atoms and the fluctuations of each node are assumed to be isotropic and Gaussian. Conformational mobility profiles in the essential space of low frequency modes were obtained using ANM server^109^ and DynOmics server.^110^

### Mutational Scanning and Sensitivity Analysis

We conducted mutational scanning analysis of the binding epitope residues for the SARS-CoV-2 S protein complexes. Each binding epitope residue was systematically mutated using all 19 substitutions and corresponding protein stability changes were computed. BeAtMuSiC approach^111–113^ was employed that is based on statistical potentials describing the pairwise inter-residue distances, backbone torsion angles and solvent accessibilities, and considers the effect of the mutation on the strength of the interactions at the interface and on the overall stability of the complex. The binding free energy of protein-protein complex can be expressed as the difference in the folding free energy of the complex and folding free energies of the two protein binding partners:

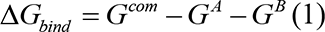

The change of the binding energy due to a mutation was calculated then as the following:

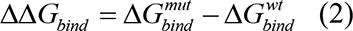

We leveraged rapid calculations based on statistical potentials to compute the ensemble-averaged binding free energy changes using equilibrium samples from MD trajectories. The binding free energy changes were computed by averaging the results over 1,000 equilibrium samples for each of the studied systems.

## SUPPORTING INFORMATION

Supporting information contains additional figures. Figure S1 shows the covariance matrixes of residue fluctuations in the S-RBD complexes with nanobodies. Figure S2 presents mutational sensitivity analysis of sites of circulating variants K417, E484, and N501 for the S-RBD complexes with Nb6 and Nb20 nanobodies. Figure S3 shows structural mapping of protein stability hotspots for the panel of studied nanobodies. Figure S4 presents mutational sensitivity analysis of sites of circulating variants for the S-RBD complexes with VHH E nanobody, biparatopic nanobody VHH VE, and CC12.3/VHH V antibody/nanobody combination.

This material is available free of charge via the Internet at http://pubs.acs.org.

## AUTHOR INFORMATION

Gennady Verkhivker, Corresponding Author

Phone: 714-516-4586; Fax: 714-532-6048;

Steve Agajanian

Deniz Yasar Oztas

Grace Gupta

The authors declare no competing financial interest.

## Acknowledgment

This work was supported by the Kay Family Foundation Grant A20-0032.

## ABBREVIATIONS

SARS: Severe Acute Respiratory Syndrome
RBD: Receptor Binding Domain
ACE2: Angiotensin-Converting Enzyme 2
NTD: N-terminal domain
RBD: receptor-binding domain.

