## Supporting Figures S1-S4 for "Atomistic Simulations and Deep Mutational Scanning of Protein Stability and Binding Interactions in the SARS-CoV-2 Spike Protein Complexes with Nanobodies: Molecular Determinants of Mutational Escape Mechanisms"

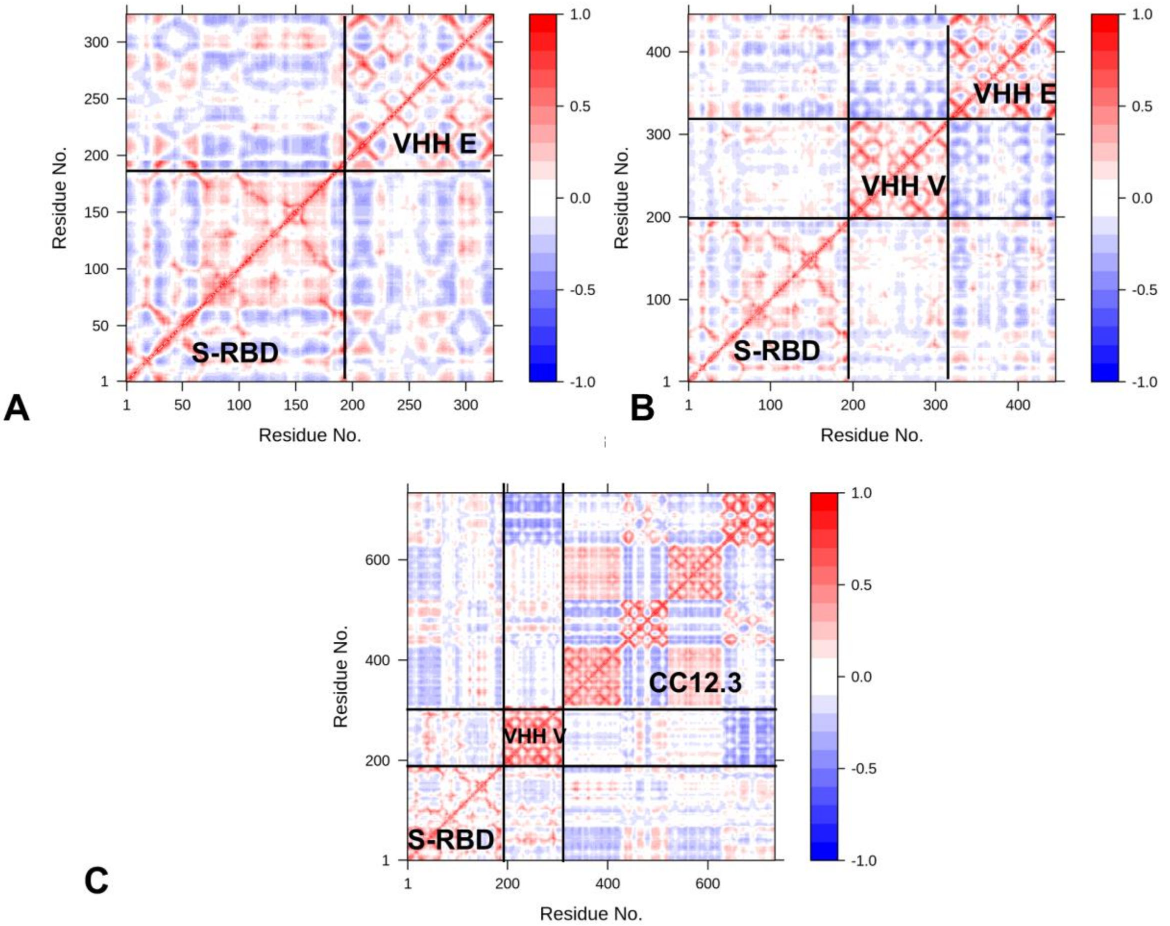


**Figure S1.** Conformational dynamics analysis and the covariance residue correlation matrixes for the SARS-CoV-2 S complexes with VHH E nanobody, pdb id 7lB14 (A), complex with the biparatopic nanobody VHH VE, pdb id 7B17 (B), and complex with CC12.3/VHH V combination, pdb id 7KN6 (C). The covariance matrix indicates coupling between pairs of residues. Cross-correlations of residue-based fluctuations vary between +1 (correlated motion; fluctuation vectors in the same direction, colored in dark red) and -1 (anti-correlated motions; fluctuation vectors in the same direction, colored in dark blue). The values > 0.5 are colored in dark red and the lower bound in the color bar indicates the value of the most anti-correlated pairs.

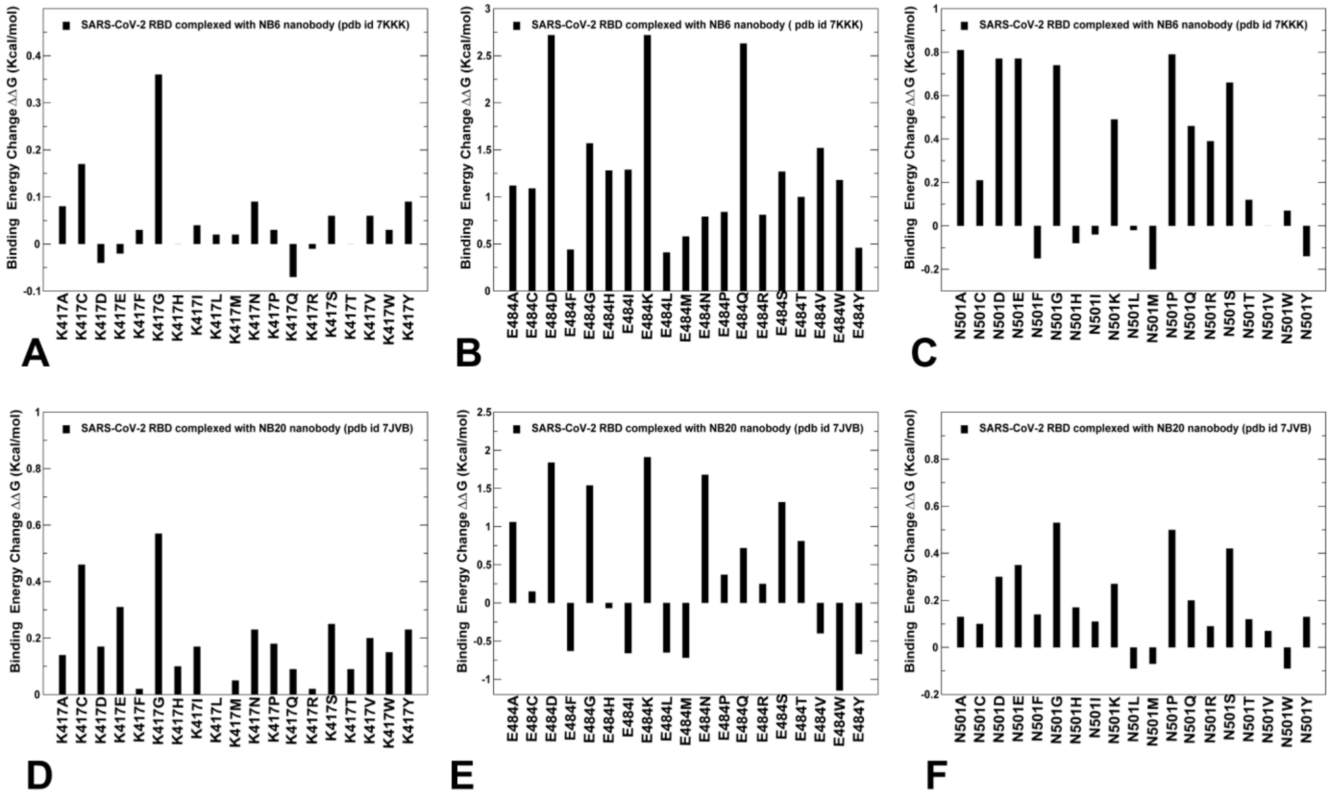


**Figure S2.** The mutational sensitivity analysis for the SARS-CoV-2 S complexes with Nb6 and Nb20 nanobodies. (A-C ) The distribution of binding free energy changes caused by mutations of K417, E484 and N501 sites in the S-RBD complexes with Nb6 nanobody. (D-F) The distribution of binding free energy changes caused by mutations of K417, E484 and N501 sites in the S-RBD complexes with Nb20 nanobody.

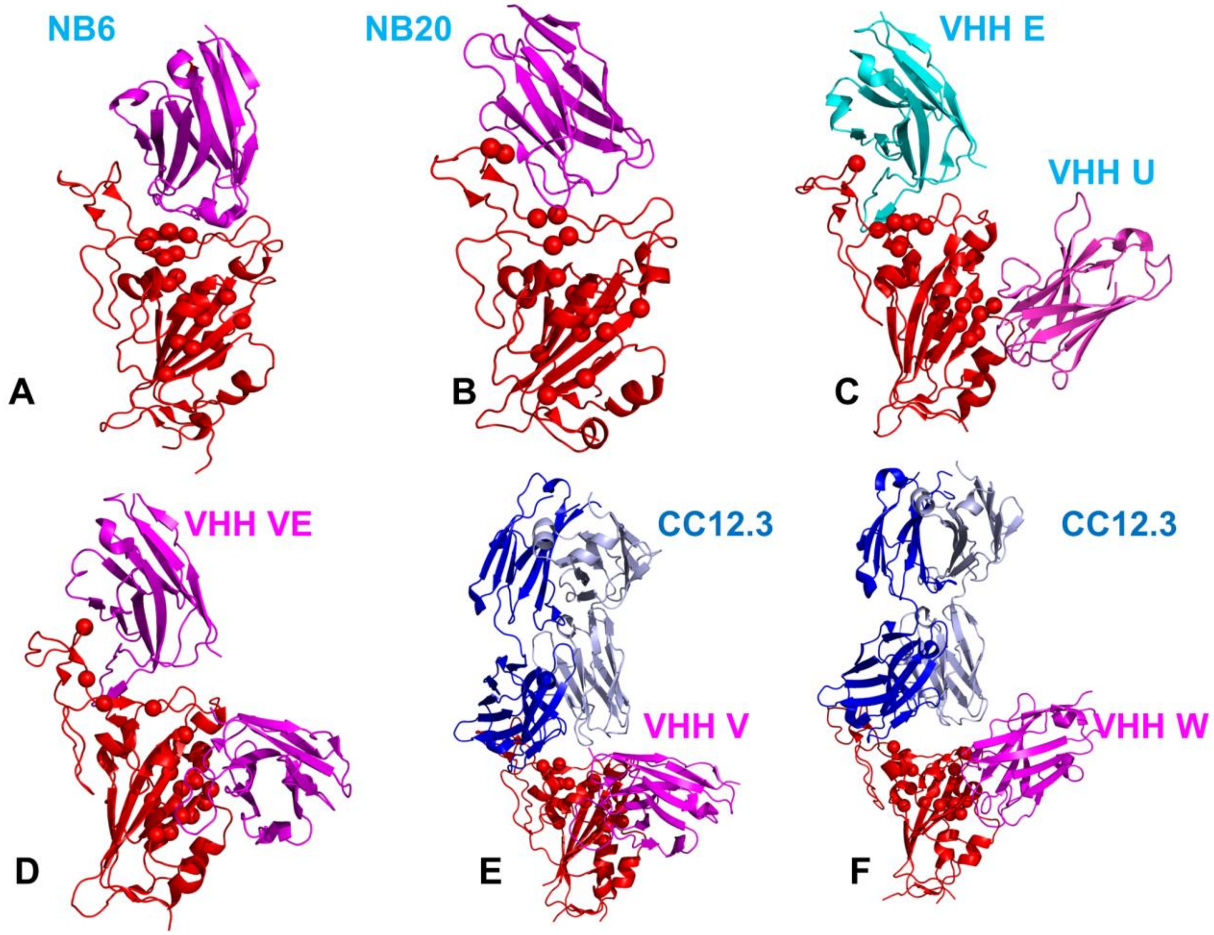


**Figure S3.** Structural mapping of protein stability centers for the SARS-CoV-2 S complexes with Nb6 (A), Nb20(B), VHH E/VHH U pair (C), biparatopic nanobody VHH VE (D), CC12.3/VHH V pair (E), and CC12.3/VHH pair (F). The S-RBD is shown in red ribbons. The bound nanobodies are shown in magenta-colored ribbons. The heavy chain of CC12.3 antibody is in blue ribbons and the light chain is in cyan-colored ribbons.

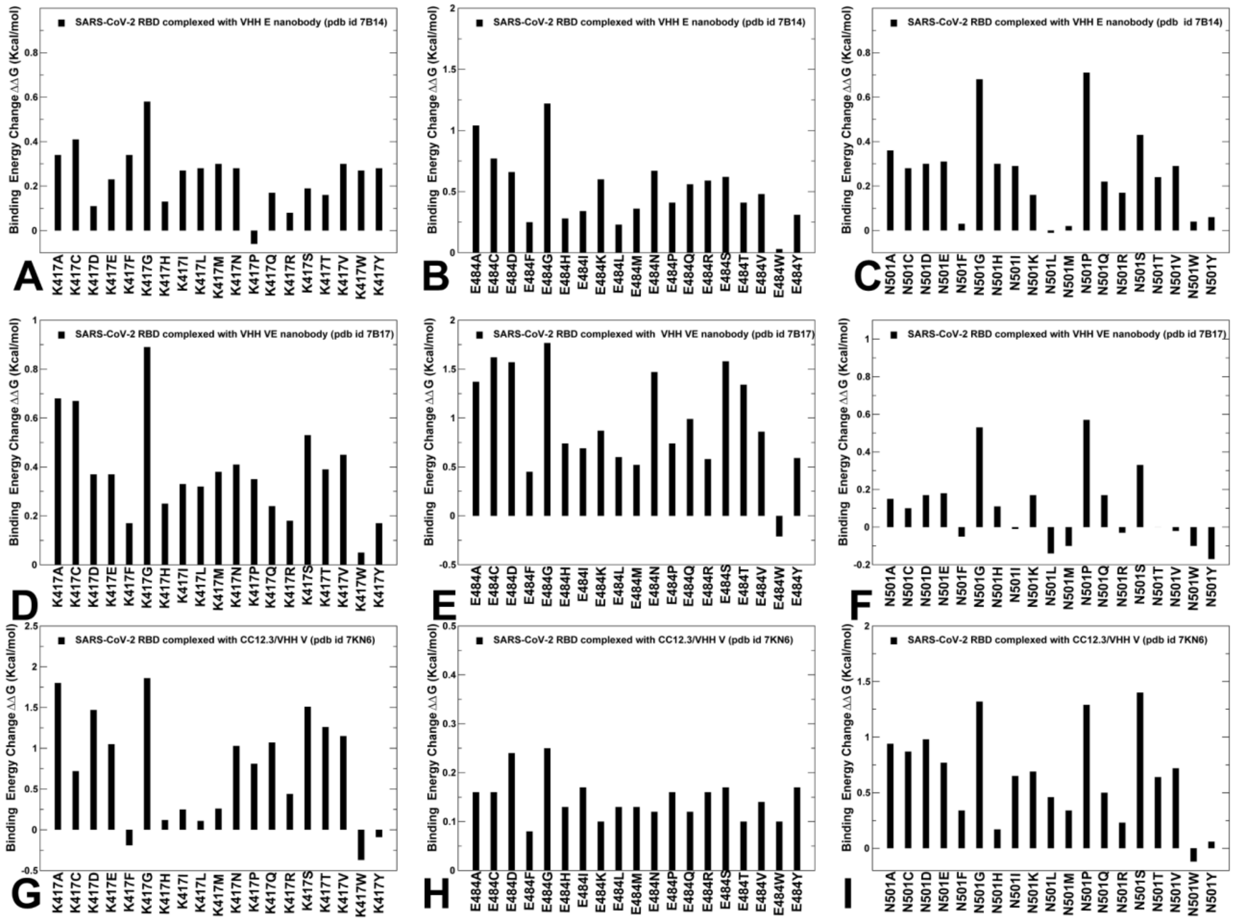


**Figure S4.** The mutational sensitivity analysis for the SARS-CoV-2 S complexes with VHH E, VHH VE nanobodies and CC12.3/VHH V antibody/nanobody combination. (A-C ) The distribution of binding free energy changes caused by mutations of K417, E484 and N501 sites in the S-RBD complexes with VHH E nanobody. (D-F) The distribution of binding free energy changes caused by mutations of K417, E484 and N501 sites in the S-RBD complexes with VHH VE biparatopic nanobody. (G-I) The distribution of binding free energy changes caused by mutations of K417, E484 and N501 sites in the S-RBD complexes with CC12.3/VHH V antibody/nanobody combination.
